# Sequence element enrichment analysis to determine the genetic basis of bacterial phenotypes

**DOI:** 10.1101/038463

**Authors:** John A. Lees, Minna Vehkala, Niko Välimäki, Simon R. Harris, Claire Chewapreecha, Nicholas J. Croucher, Pekka Marttinen, Mark R. Davies, Andrew C. Steer, Stephen Y. C. Tong, Antti Honkela, Julian Parkhill, Stephen D. Bentley, Jukka Corander

## Abstract

Bacterial genomes vary extensively in terms of both gene content and gene sequence – this plasticity hampers the use of traditional SNP-based methods for identifying all genetic associations with phenotypic variation. Here we introduce a computationally scalable and widely applicable statistical method (SEER) for the identification of sequence elements that are significantly enriched in a phenotype of interest. SEER is applicable to even tens of thousands of genomes by counting variable-length k-mers using a distributed string-mining algorithm. Robust options are provided for association analysis that also correct for the clonal population structure of bacteria. Using large collections of genomes of the major human pathogens *Streptococcus pneumoniae* and *Streptococcus pyogenes*, SEER identifies relevant previously characterised resistance determinants for several antibiotics and discovers potential novel factors related to the invasiveness of *S. pyogenes*. We thus demonstrate that our method can answer important biologically and medically relevant questions.

## Introduction

The rapidly expanding repositories of genomic data for bacteria hold an enormous and yet largely untapped potential for building a more detailed understanding of the evolutionary responses to changing environmental conditions, such as the widespread use of antibiotics and switches between host-niche as farming practices change.

Genome-wide association studies (GWAS) for bacterial phenotypes have only recently started to appear^1-5^. Use of standard GWAS methods developed originally for human SNP data have been shown to be successfully applicable to core genome mutations in bacteria^2, 3^. However, given the high level of genome plasticity of many of the known bacterial species, we can anticipate that such methods can only partially identify genetic determinants of phenotypic variation. To enable discovery of mechanisms related for instance to gene content, alternative alignment-free methods have also been introduced^1, 4^. These methods use k-mers, i.e. DNA words of length k, as generalized alternatives to SNPs as putative explanations for observed differences in phenotype distributions. The main advantage of k-mers is their ability to capture several different types of variation present across a collection of genomes, including mutations, recombinations, variable promoter architecture, differences in gene content as well as capturing these variations in regions not present in all genomes.

The previous study using k-mers to overcome limitations of SNP-based association used Monte-Carlo simulations of word gain and loss along an inferred phylogeny to control for population structure^1^, whereas SNP-based studies have used clustering algorithms on a core alignment and stratified association tests on the resulting groups of samples^2, 3^. The former does not scale computationally to the hundreds of isolates required to find lower effect-size associations, and the latter requires a core alignment, which lacks sensitivity and difficult to produce when there is a large number of samples, or they are particularly diverse.

Here we present a sequence element enrichment analysis (SEER), a method computationally scalable to tens of thousands of genomes, implemented as a stand-alone pipeline that uses either *de novo* assembled contigs or raw read data as input. We apply SEER to both simulated and real data from large and diverse populations, and show that it can accurately detect associations with antibiotic resistance caused by both presence of a gene and by SNPs in coding regions, as well as discover novel invasiveness factors.

## Results

### Implementation

SEER implements and combines three key insights which we discuss in turn: an efficient scan of all possible k-mers with a distributed string mining algorithm, an appropriate alignment-free correction for clonal population structure, and a fast and fully robust association analysis of all counted k-mers.

K-mers allow simultaneous discovery of both short genetic variants and entire genes associated with a phenotype. Longer k-mers provide higher specificity but less sensitivity than shorter k-mers. Rather than arbitrarily selecting a length prior to analysis or having to count k-mers at multiple lengths and combine the results, we provide an efficient implementation that allows counting and testing simultaneously at all k-mers at lengths over 9 bases long.

We offer three different methods to count k-mers in all samples in a study. For very large studies, or for counting directly from reads rather than assemblies, we provide an implementation of distributed string mining (DSM)^6, 7^ which limits maximum memory usage per core, but requires a large cluster to run. For data sets up to around 5 000 sample assemblies we have implemented a single core version fsm-lite (https://github.com/nvalimak/fsm-lite). For comparison with older datasets, or where resources do not allow the storage of the entire k-mer index in memory, DSK^8^ is used to count a single k-mer length in each sample individually, the results of which are then combined.

To correct for the clonal population structure of bacterial populations, a distance matrix is constructed from a random subsample of these k-mers, on which multidimensional scaling is performed (Supplementary figure 1). Compared with modelling SNP variation^9^, use of k-mers as variable sequence elements has been previously shown to accurately estimate bacterial population structure. The projections of each sample in three dimensions are used as covariates to control for the clonal population structure. Simulations of bacterial genomes using a known tree showed this method gave a higher resolution control than using only population clustering (Supplementary figure 2). Before testing for association we filter k-mers based on their frequency and unadjusted p-value to reduce false positives from testing underpowered k-mers and reduce computational time.

Then, for each k-mer, a logistic curve is fitted to binary phenotype data, and a linear model to continuous data, using a time efficient optimisation routine to allow testing of all k-mers. Bacteria can be subject to extremely strong selection pressures, producing common variants with very large effect sizes, such as antibiotics inducing resistance-conferring variants. This can make the data perfectly separable, and consequently the maximum likelihood estimate ceases to exist for the logistic model. Firth regression^10^ has been used to obtain results in these cases.

For the basal cut-off for significance we use *p* < 0.05, which in our testing we conservatively Bonferroni corrected to the threshold 1x10^-8^ based on every position in the S. pneumoniae genome having three possible mutations^11^, and all this variation being uncorrelated. This is a strict cut-off level that prevents a large number of false-positives due to the extensive amount of k-mers being tested, but does not over-penalise by correcting directly on the basis of the number of k-mers counted. Simulations suggested a cut-off of 1.4x10^-8^ would be appropriate, supporting this reasoning. Association effect size and p-value of the MDS components can also be included in the output, to compare lineage and variant effects on the phenotype variation.

K-mers reaching significance are filtered post-association and mapped onto both a well-annotated reference sequence and the annotated draft assemblies to allow discovery of variation in accessory genes not present in the reference strain. The significant k-mers themselves can also be assembled into a longer consensus sequence. Annotating variants by predicted function and effect (against a reference sequence) in the resulting k-mers facilitates fine-mapping of SNPs and small indels.

Meta-analysis of association studies increases sample size, which improves power and reduces false-positive rates^12^. To facilitate meta-analysis of k-mers across studies, the output of SEER includes effect size, direction and standard error, which can be used directly with existing software to meta-analyse all overlapping k-mers.

SEER is implemented in C++, and available at https://github.com/johnlees/seer as source code and a pre-compiled binary.

### Application to simulated data

To test the power of SEER across different sample sizes, we simulated 3 069 *Streptococcus pneumoniae* genomes from the phylogeny observed in a Thai refugee camp^13^ using parameters estimated from real data including accumulation of SNPs, indels (Supplementary figure 3), gene loss and recombination events. Using knowledge of the true alignments, we then artificially associated an accessory gene with a phenotype over a range of odds-ratios and evaluated power at different sample sizes (Fig. 1a). The expected pattern for this power calculation is seen, with higher odds-ratio effects being easier to detect. Currently detected associations in bacteria have had large effect sizes (OR > 28 host-specificity^1^, OR > 3 beta-lactam resistance^2^), and the required sample sizes predicted here are consistent with these discoveries.

**Fig. 1:**
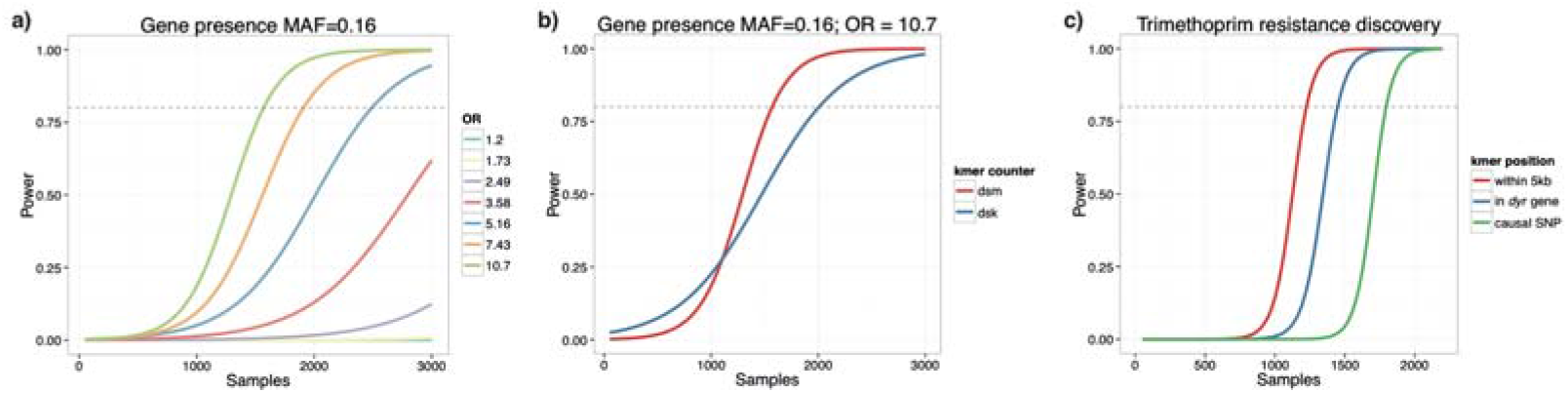
Using simulations and subsamples of the population as described in the online methods, power for a) detecting gene presence/absence at different oddsratios b) using all informative k-mers versus a single length c) detecting k-mers near, in the correct gene, or containing the causal variant for trimethoprim resistance. All curves are logistic fits to the mean power over 100 subsamples.

The large k-mer diversity, along with the population stratification of gene loss, makes the simulated estimate of the sample size required to reach the stated power clearly conservative. Convergent evolution along multiple branches of a phylogeny for a real population reacting to selection pressures will reduce the required sample size^14^.

We also used k-mers counted at constant lengths by DSK to perform the gene presence/absence association (Fig. 1b). Counting all informative k-mers rather than a range of pre-defined k-mer lengths gives greater power to detect associations, with 80% power being reached at around 1 500 samples, compared with 2 000 samples required by the pre-defined lengths. The slightly lower power at low sample numbers is due to a stricter Bonferroni adjustment being applied to the larger number of DSM k-mers over the DSK k-mers. This is exactly the expected advantage from including shorter k-mers to increase sensitivity, but as k-mers are correlated with each other due to evolving along the same phylogeny, using the same Bonferroni correction for multiple testing does not decrease specificity.

The strong linkage disequilibrium (LD) caused by the clonal reproduction of bacterial populations means that non-causal k-mers may also appear to be associated. This is well documented in human genetics; non-causal variants tag the causal variant increasing discovery power, but make it more difficult to fine-map the true link between genotype and phenotype^15^. In simulations it is difficult to replicate the LD patterns observed in real populations, as recombination maps for specific bacterial lineages are not yet known. To evaluate fine-mapping power of a SNP we instead used the real sequence data and simulated phenotypes based on changing the effect size of a known causal variant and evaluating the physical distance of significant k-mers from the variant site.

Using DSM we counted 68M k-mers which we then tested for association. The 2 639 significant k-mers were placed into three categories if after mapping to a reference genome they contained the causal variant I100L (10), were within the same gene (74), or within 2.5kb in either direction (207). Figure 1c) shows the resulting power when random subsamples of the population are taken. As expected, power is higher when not specifying that the causal variant must be hit, as there are many more k-mers which are in LD with the SNP than directly overlapping it, thus increasing sensitivity.

### Confirmation of known resistance mechanisms in a large population of *S. pneumoniae*

SEER was applied to the sequenced genomes from the study described above, using measured resistance to five different antibiotics as the phenotype: chloramphenicol, erythromycin, β-lactams, tetracycline and trimethoprim. Chloramphenicol resistance is conferred by the *cat* gene on the integrative conjugative element (ICE) Tn*5253* in the *S. pneumoniae* chromosome, and similarly tetracycline resistance is conferred by the *tetM* gene which is also carried on the ICE^16^. For both of these drug resistance phenotypes the ICE contains 99% of the significant k-mers, and the causal genes rank highly within the clusters (Table 1, Supplementary figure 4).

**Table 1:** Results from SEER for antibiotic resistance binary outcome on a population of 3069 *S. pneumoniae*. Significant k-mers are first interpreted by mapping to the ATCC 700669 reference genome. Up to the first four highest covered annotations are shown, and if the known mechanism is amongst these it is highlighted in orange. The ICE is the top hit in three analyses, as it carries multiple drug-resistance elements and is commonly found in multi-drug resistant strains^16^. The distribution of phenotype across the phylogeny is shown in Supplementary figure 5.

| Antibiotic | Resistant samples | Number of significant k-mers |  |  |  |
| --- | --- | --- | --- | --- | --- |
|  |  | Total | Mapped to reference | Highest coverage annotation | Causal element |
| Chloramphenicol | 204 (7%) | 1 526 | 1 526 | 1 508 – ICE<br>288 – ORF (UniParc B8ZK82)<br>206 – rep<br>166 – cat | 166 – cat |
| Erythromycin | 803 (26%) | 1 154 | 112 | 10 – permease (UniParc B8ZKV5)<br>8 – prfC<br>6 – gatA<br>4 – ICE | 4 – mega element<br>2 – mef<br>2 – omega element |
| β-lactams | 1 563 (51%) | 23 876 | 17 453 | 381 – ICE<br>145 – prophage MM1<br>50 – SPN23F15110 (UniParc B8ZLE7)<br>49 – ICE orf16 | 47 – pbp2x<br>20 – pbp2b<br>8 – pbp1a |
| Tetracycline | 1 958 (64%) | 962 | 962 | 962 – ICE<br>136 – ICE orf16<br>121 – ICE orf15<br>96 – tetM | 96 – tetM |
| Trimethoprim | 2 553 (83%) | 2 639 | 210 | 21 – dyr | 21 – dyr |

Resistance to erythromycin is also conferred by presence of a gene, but there are multiple genes that can perform the same function (*ermB*, *mef*, mel)^17^. In the population studied, this phenotype was strongly associated with two large lineages (Supplementary figure 5), making the task of disentangling association with a lineage versus a specific locus more difficult. Significant k-mers are found in the mega and omega cassettes, which carry the *mel/mef* and *ermB* resistance elements respectively. Some k-mers do not map to the reference, as they are due to lineage specific associations with genetic elements not found in the reference strain. This highlights both the need to map to a close reference or draft assembly to interpret hits, as well as the use of functional follow-up to validate potential hits from SEER.

Multiple mechanisms of resistance to β-lactams are possible^2^. Here, we consider just the most important (i.e. highest effect size) mutations, which are SNPs in the penicillin binding proteins *pbp2x, pbp2b* and *pbp1a*. In this case looking at highest coverage annotations finds these genes, but is not sufficient as so many k-mers are significant – either due to other mechanisms of resistance, physical linkage with causal variants or co-selection for resistance conferring mutations. Instead, looking at the k-mers with the most significant p-values gives the top four hit loci as *pbp2b* (p=10^-132^), *pbp2x* (p=10^-96^), putative RNA pseudouridylate synthase UniParc B8ZPU5 (p=10^-92^) and *pbp1a* (p=10^-89^). The non-*pbp* hit is a homologue of a gene in linkage disequilibrium with *pbp2b*, which would suggest mismapping rather than causation of resistance.

Trimethoprim resistance in *S. pneumoniae* is conferred by the SNP I100L in the *folA/dyr* gene^18^. The *dpr* and *dyr* genes, which are adjacent in the genome, have the highest coverage of significant k-mers (Fig. 2). Following our fine-mapping procedure, we call four high-confidence SNPs that are predicted to be more likely to affect protein function than synonymous SNPs. One is the causal SNP, and the others appear to be hitchhikers in LD with I100L. By evaluating whether sites are conserved across the protein family^19^, the known causal SNP is ranked as the highest variant, showing that in this case fine-mapping is possible using the output from SEER.

**Fig. 2:**
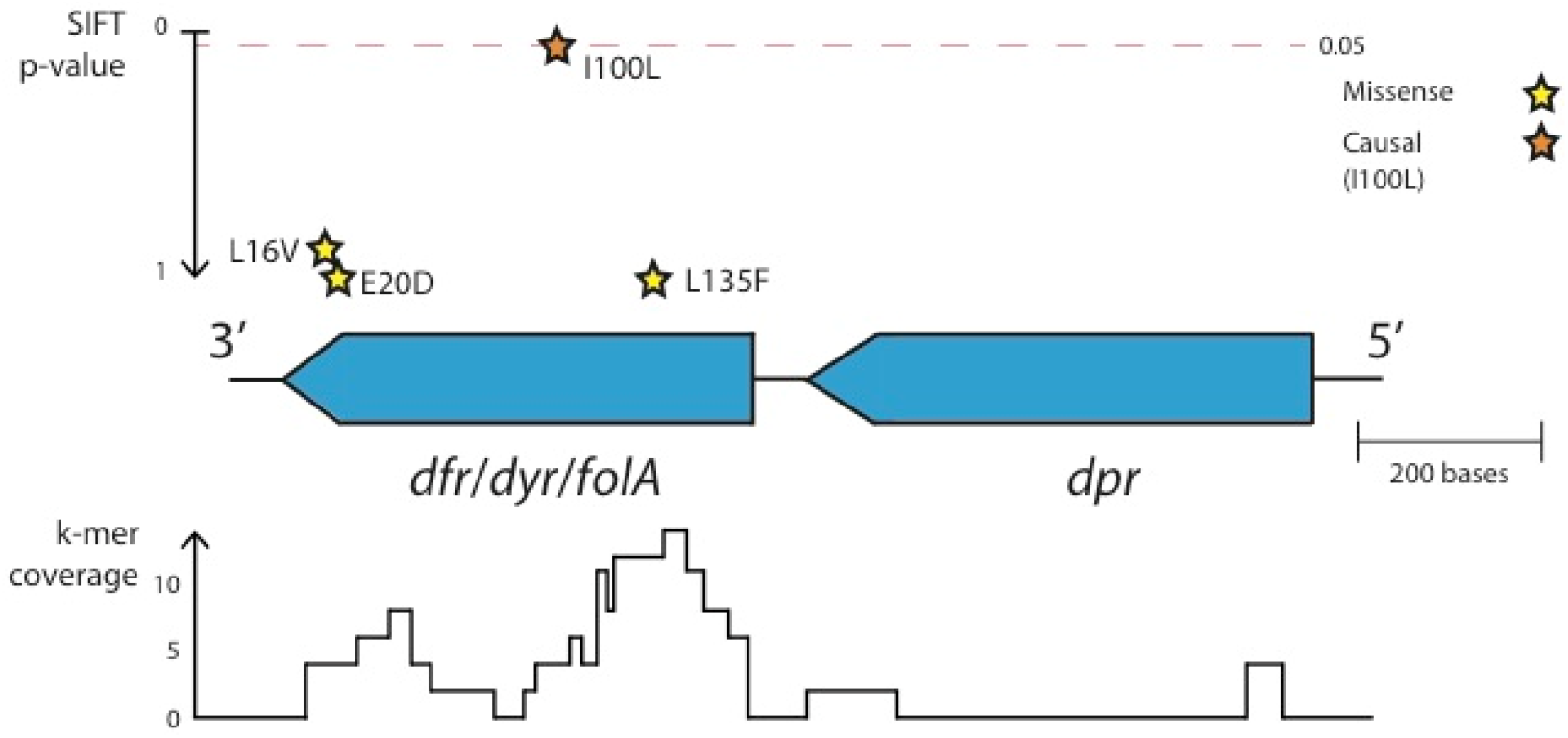
Fine mapping trimethoprim resistance. The locus pictured contains 72 significant k-mers, the most of any gene cluster. Coverage over the locus is pictured at the bottom of the figure. Shown above the genes are high quality missense SNPs, plotted using their p-value for affecting protein function as predicted by SIFT.

We then compared the results from SEER with the results from two existing methods (as described in online methods). The first method uses mapping of SNPs against a reference, followed by applying the Cochran-Mantel-Haenszel test at every variable site^2^. The second uses dsk^8^ to count k-mers of length 31, and a highly robust correction for population structure which scales to around 100 genomes^1^.

The results are shown in supplementary table 1. Both SEER and association of a core mapping of SNPs identify resistances caused by presence of a gene, when it is present in the reference used for mapping. Both produce their most significant p-values in the causal element, though SEER appears to have a lower false-positive rate. However, as demonstrated by chloramphenicol resistance, if not enough SNP calls are made in the causal gene this hinders fine-mapping. SNP-mediated resistance showed the same pattern since many other SNPs were ranked above the causal variant. In the case of β-lactam resistance both methods seem to perform equally well, likely due to the higher rate of recombination and the creation of mosaic *pbp* genes.

Additionally, as for erythromycin resistance, when an element is not present in the reference SNPs have been called against it is not detectable in SNP-based association analysis. In such cases multiple mappings against other reference genomes would have to be made, which is a tedious and computationally costly procedure. Alternatively a draft assembly with the phenotype from the study could be picked as a second reference to map to, however this may be lower quality than those in public databases picked by genetic content rather than phenotype, and would not necessarily be able to detect multiple genetic mechanisms (as in the case of erythromycin resistance, no single sequenced genome contains all known resistance mechanisms).

Since the k-mer results from SEER are reference-free, these issues are avoided as just the significant k-mers can quickly be mapped to all available references. Alternatively, the significant k-mers can be mapped to all draft assemblies in the study, at least one of which is guaranteed to contain the k-mer, to check if any annotations are overlapped.

For the small sample, 31mer approach significance was not reached for chloramphenicol, tetracycline or trimethoprim as the effect size of any k-mer is too small to be detected in the number of samples accessible by the method. Erythromycin had 19 307 hits, and β-lactams 419 hits, at between 1-2% MAF which are all false positives that would likely have been excluded by a fully robust population structure correction method.

### Discovery of conjugative elements associated with *Streptococcus pyogenes* isolation location and invasiveness

Most bacterial GWAS studies to date have searched for genotypic variants that contribute towards or completely explain antibiotic resistance phenotypes. As a proof of principle that SEER can be used for the discovery stage of sequence elements associated with other clinically important phenotypes, we applied our tool to 675 *S. pyogenes* (group A *Streptococcus*) genomes from invasive and non-invasive isolates.

The top hit was the *tetM* gene in a conjugative transposon (Tn*916*) carried by 23% of isolates (Supplementary figures 6 and 7). These elements are variably present in the chromosome of *S. pyogenes^20^*, and the lack of co-segregation with population structure explains our power to discover the association. However, as a different proportion of the isolates from each collection were invasive (Fiji - 13%; Kilif - 43%), the significant k-mers will also include elements specific to Kilifi. Indeed, we found that this version of Tn916 was never present in genomes collected from Fiji. When country of isolation was included as a covariate in the regression, these hits were no longer significant – highlighting the importance of such considerations in performing association studies in large bacterial populations.

After applying this correction, we find two significant hits (Supplementary figure 8). The first corresponds to SNPs associating a specific allele of *pepF* (Oligoendopeptidase F; UniProt:P54124) with invasive isolates. This could indicate a recombination event, due to the high SNP density and discordance with vertical evolution with respect to the inferred phylogeny^21, 22^. The second hit represents SNPs in the intergenic region upstream of both IgG-binding protein H and *nrdl* (ribonucleotide reductase). If this were found to affect expression of the IgG-binding protein, this would be a plausible novel genetic mechanism affecting pathogenesis^23, 24^.

The association of both of these variations would have to be validated either *in vitro* or a replication cohort, and functional follow-up such as RNA-seq may also further help with their interpretation.

Applying a Cochran-Mantel-Haenszel test to SNPs called against a reference sequence found no sites significantly associated with invasiveness. The *tetM* gene and transposon are not found in the reference sequence, and therefore cannot be discovered by this method. The population structure is so diverse that 88 different clusters are found, which overcorrects leaving too few samples within each group to have power to discover associations.

## Discussion

SEER is a reference-independent, scalable pipeline capable of finding bacterial sequence elements associated with a range of phenotypes while controlling for clonal population structure. The sequence elements can be interpreted in terms of protein function using sequence databases, and we have shown that even single causal variants can be fine-mapped using the SEER output.

Our use of all informative k-mers together with robust regression methods, and the ability to analyse very large sample sizes show improved sensitivity over existing methods. This provides a generic approach capable of analysing the rapidly increasing number of bacterial whole genome sequences linked with a range of different phenotypes. The output can readily be used in a meta-analysis of sequence elements to facilitate the combination of new studies with published data, increasing both discovery power and confirming the significance of results. As with all association methods, our approach is limited by the amount of recombination and convergent evolution that occurs in the observed population, since the discovery of causal sequence elements is principally constrained by the extent of linkage disequilibrium. However, by introducing improved computational scalability and statistical sensitivity SEER significantly pushes the existing boundaries for answering important biologically and medically relevant questions.

## Online methods

### Counting informative k-mers in samples

Over all *N* samples, all k-mers over 9 bases long that occur in more than one sample are counted. All non-informative k-mers are omitted from the output; a k-mer *X* is not informative if any one base extension to the left (*aX*) or right (*Xa*) has exactly the same frequency support vector as *X*. The frequency support vector has *N* entries, each being the number of occurrences of k-mer *X* in that sample. Further filtering conditions are explained in the sections below.

Distributed string mining (DSM)^6, 7^ parallelises to as much as one sample per core, and either 16 or 64 master server processes. DSM includes an optional entropy-filtering setting that filters the output k-mers based on both number of samples present and frequency distribution. On our 3 069 simulated genomes this took 2 hrs 38 min on 16 cores, and used 1Gb RAM. The distributed approach is applicable up to terabytes of short-read data^7^, but requires a cluster environment to run. As an easy-to-use alternative, we propose a single core version of DSM that is applicable for gigabyte-scale data. We implemented the single core version based on a succinct data structure library^25^ to produce the same output as DSM. On 675 S. pyogenes genomes this took 3hrs 44min and used 22.3Gb RAM.

To count single k-mer lengths, an associative array was used to combine the results from DSK in memory. We concatenated results from k-mer lengths of 21, 31 and 41, as in previous studies^1^. This can scale to large genome numbers by instead using external sorting to avoid storing the entire array in memory.

### Filtering k-mers

K-mers are filtered if either they appear in <1% or >99% of samples, or are over 100 bases long. We also test if the p-value of association in a simple *χ*^2^ test (1 d.f.) is less than 10^-5^, as in simulations this was true for all true positives. In the case of a continuous phenotype a Welch two-sample t-test is used instead.

### Covariates to control for population structure

A random sample of between 0.1% and 1% of k-mers appearing in between 5-95% of isolates is taken. We then construct a pairwise distance matrix **D**, with each element being equal to a sum over all *m* sampled k-mers:

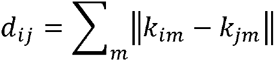

where *k_im_* is 1 if the *m*th sampled k-mer is present in sample *i*, and 0 otherwise.

Metric multi-dimensional scaling is then performed, projecting these distances into three dimensions. The normalised eigenvectors of each dimension are used as covariates in the regression model. The number of dimensions used is a user-adjustable parameter, and can be evaluated by the goodness-of-fit and the magnitude of the eigenvalues. In species tree with two lineages and 96 isolates one dimension was sufficient as a population control, whereas for the larger collection of 3069 isolates 10-15 dimensions were needed to give tight control (Supplementary figure 9). Over all our studies, generally three dimensions appeared a good trade-off between sensitivity and specificity.

### Logistic and linear regression

For samples with binary outcome vector *y*, for each k-mer a logistic model is fitted:

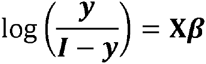

where absence and presence for each k-mer coded as 0 and 1 respectively in column 2 of the design matrix **X** (column 1 is a vector of ones, giving an intercept term). Subsequent columns *j* of **X** contain the eigenvectors of the MDS projection, user-supplied categorical covariates (dummy encoded), and quantitative covariates (normalised). The BFGS algorithm is used to maximise the log likelihood *L* in terms of the gradient vector ***β*** (using an analytic expression for d(log *L*)d**β**):

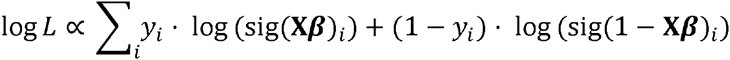

where sig is the sigmoid function. If this fails to converge, *n* Newton-Raphson iterations are applied to ***β***

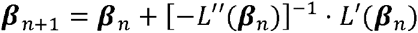

from a starting point using the mean phenotype as the intercept, and the root-mean squared beta from a test of k-mers passing filtering

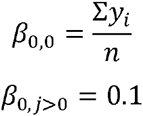

which is slower, but has a higher success rate. If this fails to converge due to the observed points being separable in the high dimensional space, or the standard error of the slope is greater than 3 (which empirically indicated almost separable data, with no counts in one element of the contingency table), Firth logistic regression^10^ is then applied. This adds an adjustment to log *L*:

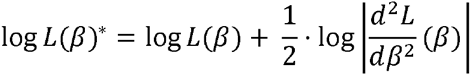

using which Newton-Raphson iterations are applied as above.

In the case of a continuous phenotype a linear model is fitted:

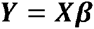

The squared distance U(**β**)

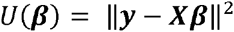

is minimised using the BFGS algorithm. If this fails to converge then the analytic solution is obtained by orthogonal decomposition:

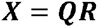

then back-solving for **β** in:

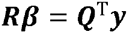

In both cases the standard error on β_1_ is calculated by inverting the Fisher information matrix d^2^L/d**β**^2^ (inversions are performed by Cholesky decomposition, or if this fails due to the matrix being almost singular the Moore-Penrose pseudoinverse is taken) to obtain the variance-covariance matrix. The Wald statistic is calculated with the null hypothesis of no association (β_1_ = 0) :

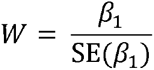

which is the test statistic of a χ^2^ distribution with 1 d.f. This is equivalent to the positive tail of a standard normal distribution, the integral of which gives the p-value. To calculate an empirical significance testing cut-off for the p-value under multiple correlated tests, we observed the distribution of p-values from 100 random permutations of phenotype. Setting the family-wise error rate (FWER) at 0.05 gave a cut-off of 1.4×10^-8^.

### SEER implementation

SEER is implemented in C++ using the armadillo linear algebra library^26^, and dlib optimisation library^27^. On a simulation of 3 069 diverse 0.4Mb genomes, 143M k-mers were counted by DSM and 25M 31-mers by DSK. On the largest DSM set, using 16 cores and subsampling 300 000 k-mers (0.2% of the total), calculating population covariates took 6hr 42min and 8.33GB RAM. This step is O(N^2^M) where N is number of samples and M is number of k-mers, but can be parallelised across up to N^2^ cores.

Processing all 143M informative k-mers as described took 69min 44s and 23MB RAM on 16 cores. This step is O(M) and can be parallelised across up to M cores.

On the real dataset of full length genomes the 68M informative k-mers counted was less than the simulated dataset above, as the parameters of the simulation created particularly diverse final genomes.

### Interpreting significant k-mers

K-mers reaching the threshold for significance are then post-association filtered requiring β_1_ > 0 as a negative effect size does not make biological sense. Remaining k-mers are searched for by exact match in their *de novo* assemblies, and annotations of features examined for overlap of function. BLAT^28^ is also used with a step size of 2 and minimum match size of 15 to find inexact but close matches to a well annotated reference sequence.

To better search for gene clusters associated with phenotype, these k-mers are assembled using Velvet^29^ choosing a smaller sub-k-mer size which maximises longest contig length of the final assembly. K-mers which are then substrings of others significant k-mers are removed.

### Mapping of a single SNP

Using the BLAT mapping of significant k-mers to a reference sequence, SNPs are called using bcftools^30^. Quality scores for a read are set to be identical, and are set as the Phred-scaled Holm-adjusted p-values from association. High quality (QUAL > 100) SNPs are then annotated for function using SnpEff^31^, and the effect of missense SNPs on protein function is ranked using SIFT^19^.

### Comparison to existing methods

We compare to two existing methods. The first uses a core-genome SNP mapping along with population clusters defined from the same alignment to perform a Cochran-Mantel-Haenszel test at every called variant site^2^. The second uses a fixed k-mer length of 31 as counted by dsk^8^, with a Monte Carlo phylogeny-based population control^1^. As the second method is not scalable to this population size we used our population control as calculated from all genomes in the population, and a subsample of 100 samples to calculate association statistics, which is roughly the number computationally accessible by this method. In both cases, the same Bonferroni correction is used as for SEER.

### Simulating bacterial populations

A random subset of 450 genes from the *Streptococcus pneumoniae* ATCC 700669^16^ strain were used as the starting genome for ALF^32^. ALF simulated 3069 final genomes along the phylogeny observed in a Thai refugee camp^13^. An alignment between *S. pneumoniae* strains R6, 19F and *Streptococcus rnitis* B6 using Progressive Cactus was used to estimate rates in the GTR matrix and the size distribution of insertions and deletions (INDELs – Supplementary figure 3). Previous estimates for the relative rate of SNPs to INDELs^33^ and the rate of horizontal gene transfer and loss^13^ were used.

pIRS^34^ was used to simulate error-prone reads from genomes at the tips of the tree, which were then assembled by Velvet^29^. DSM was used to count k-mers from these *de novo* assemblies.

To test the similarity of the population control to existing methods, 96 full *Streptococcus pneumoniae* ATCC 700669 genomes were evolved with ALF. Intergenic regions were also evolved using Dawg^35^ at a previously determined rate^36^. These were combined, and assemblies generated and k-mers counted as above. A distance matrix was created from 1% of the k-mers as described above, and a neighbour-joining tree produced from this.

The resulting tree was ranked against the true tree by counting one for each pair of isolates in each BAPS^37^ cluster which had an isolate not in the same BAPS cluster as a descendent of their MRCA.

### Simulating phenotype based on genotype and odds-ratio

Ratio of cases to controls in the population (*S_R_*) was set at 50% to represent antibiotic resistance, and a single variant (gene presence/absence or a SNP) was designated as causal. Minor allele frequency (MAF) in the population is set from the simulation, and odds-ratio (OR) can be varied. The number of disease cases *D_E_* is then the solution to a quadratic equation^38^, which is related to probability of a sample being a case by:

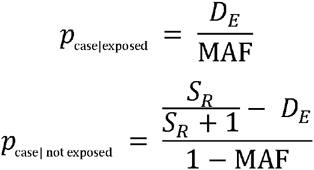

The population was then randomly subsampled 100 times, with case and control status assigned for each run using these formulae. Power was defined by the proportion of runs that had at least one k-mer in the gene associated with phenotype reaching significance.

### Elements enriched in *S. pyogenes* invasiveness

We sequenced 675 isolates of *S. pyogenes* on the Illumina HiSeq platform, of which 347 were from Fiji and 328 were from Kilifi^39^. We defined those isolated from blood, cerebrospinal fluid (CSF) or broncho-pulmonary aspirate as invasive (n = 185), and those isolated from throat, skin or urine as non-invasive (n = 490). Including country as a categorical covariate was necessary, as without doing so many elements which stratify by isolate collection appear as significant. The SEER pipeline was run as described, yielding 1233 k-mers which exceeded the threshold for significance.

BLAST of the k-mers with the nr/nt database was used to determine a suitable reference to map to, and after mapping SNPs were called as above.

## Acknowledgements

We would like to thank James Hadfield for his help in integrating SEER’s output into the bacterial genome visualisation tool JSCandy, and Jeff Barrett and his group for helpful discussions on the relation of association studies in human genetics to prokaryotic genetics.

This work was supported by Wellcome Trust grant 098051, MRC grant 1365620, ERC grant 239784, Academy of Finland grant 287665 and COIN Centre of Excellence.

## Author Contributions

JAL – Designed method, performed analysis, wrote manuscript.

MV – Designed method, performed analysis, wrote manuscript.

NV – Participated in method design, edited manuscript.

SRH – Helped with interpretation of *S. pyogenes* data

CC – Prepared genetic and metadata from Maela isolates.

NJC – Helped with interpretation of antibiotic resistance elements, edited manuscript.

PM – Participated in method design, edited manuscript.

AH – Participated in method design, edited manuscript.

JP – Advised on microbiological interpretation, edited manuscript.

SDB – Advised on microbiological interpretation, edited manuscript.

JC – Designed method, performed analysis, wrote manuscript.

## Data Access

SEER is available at https://github.com/johnlees/seer, DSM at https://github.com/HIITMetagenomics/dsm-framework and fsm-lite at https://github.com/nvalimak/fsm-lite.

Scripts used to perform the simulations are available at https://github.com/johnlees/bioinformatics

Results from the *S. pyogenes* invasiveness GWAS can be found at: http://dx.doi.org/10.6084/m9.figshare.1613851 and can be loaded directly into JSCandy (http://jameshadfield.github.io/JScandy/) to view the results.

## Supplementary data

**Supplementary table 1:**
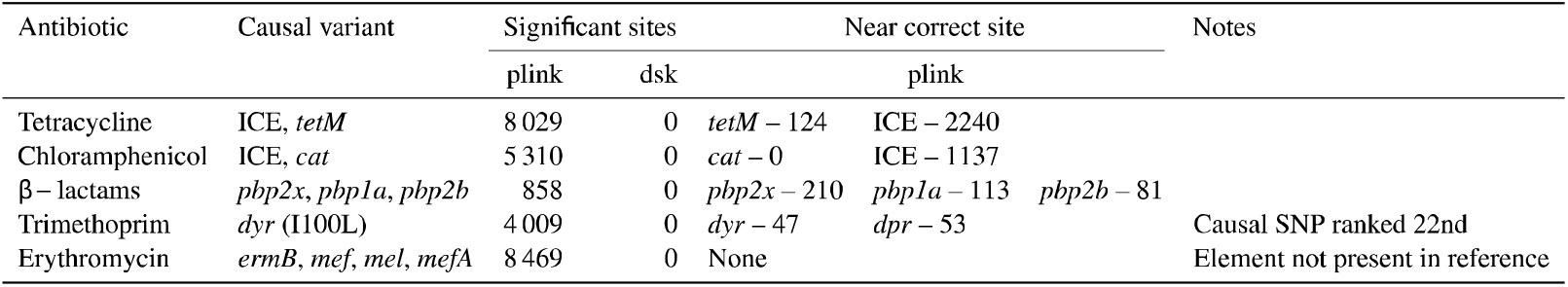
Comparison of SEER with results from existing methods in finding genetic associations with antibiotic resistance in the Chewapreecha *et. al*. study of 3069 Thai carriage *S. pneumoniae* samples. For each of the five antibiotics, the true causal variant is listed, as are the number of hits passing the significance threshold for each method (plink and dsk) and the number which map to the correct region.

**Supplementary figure 1:**
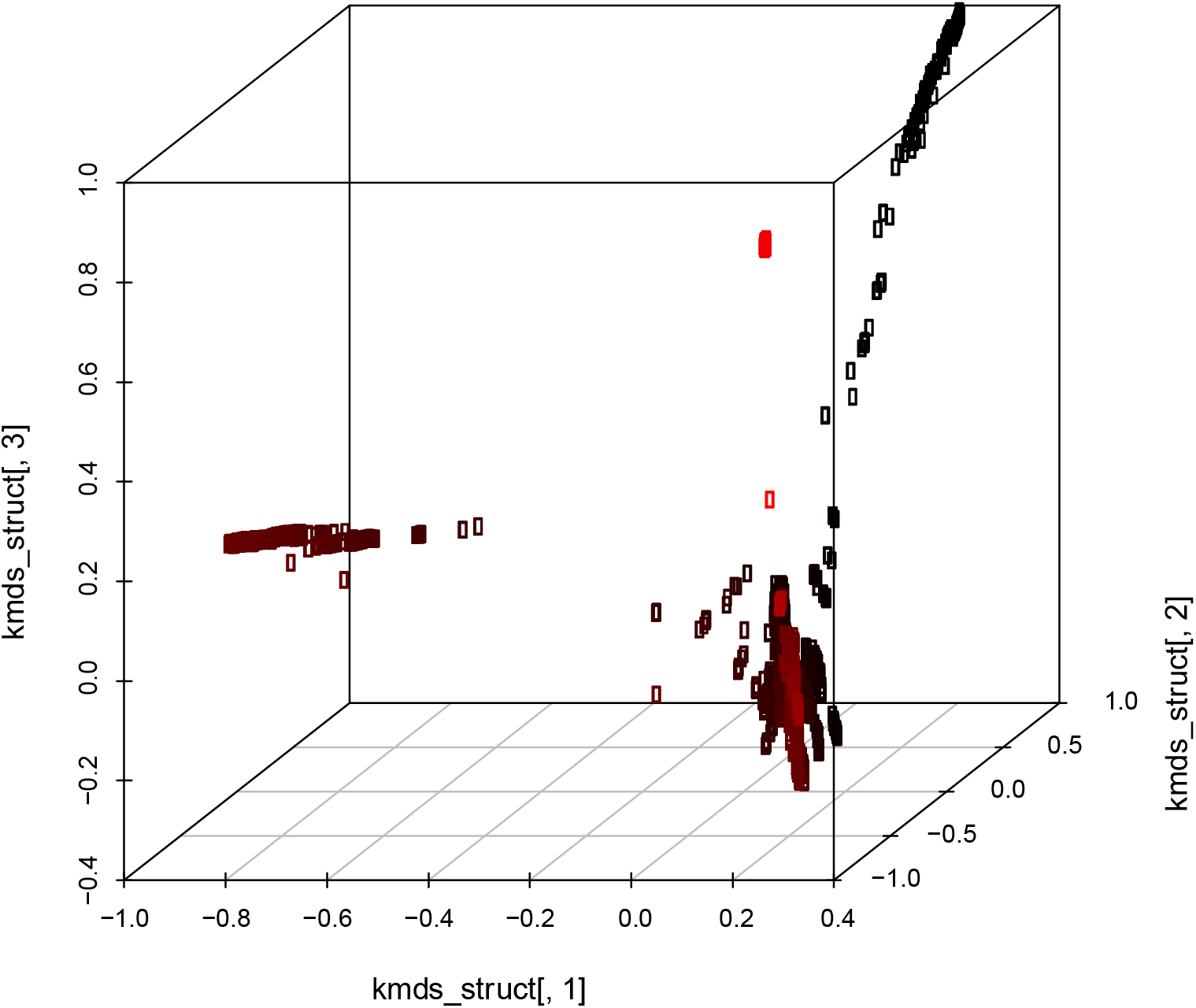
Plot of the k-mer distances projected into three dimensions by MDS for the Chewapreecha *et. al*. study of 3069 Thai carriage *S. pneumoniae* samples. Shade from black to red is by y-coordinate (2^nd^ MDS component).

**Supplementary figure 2:**
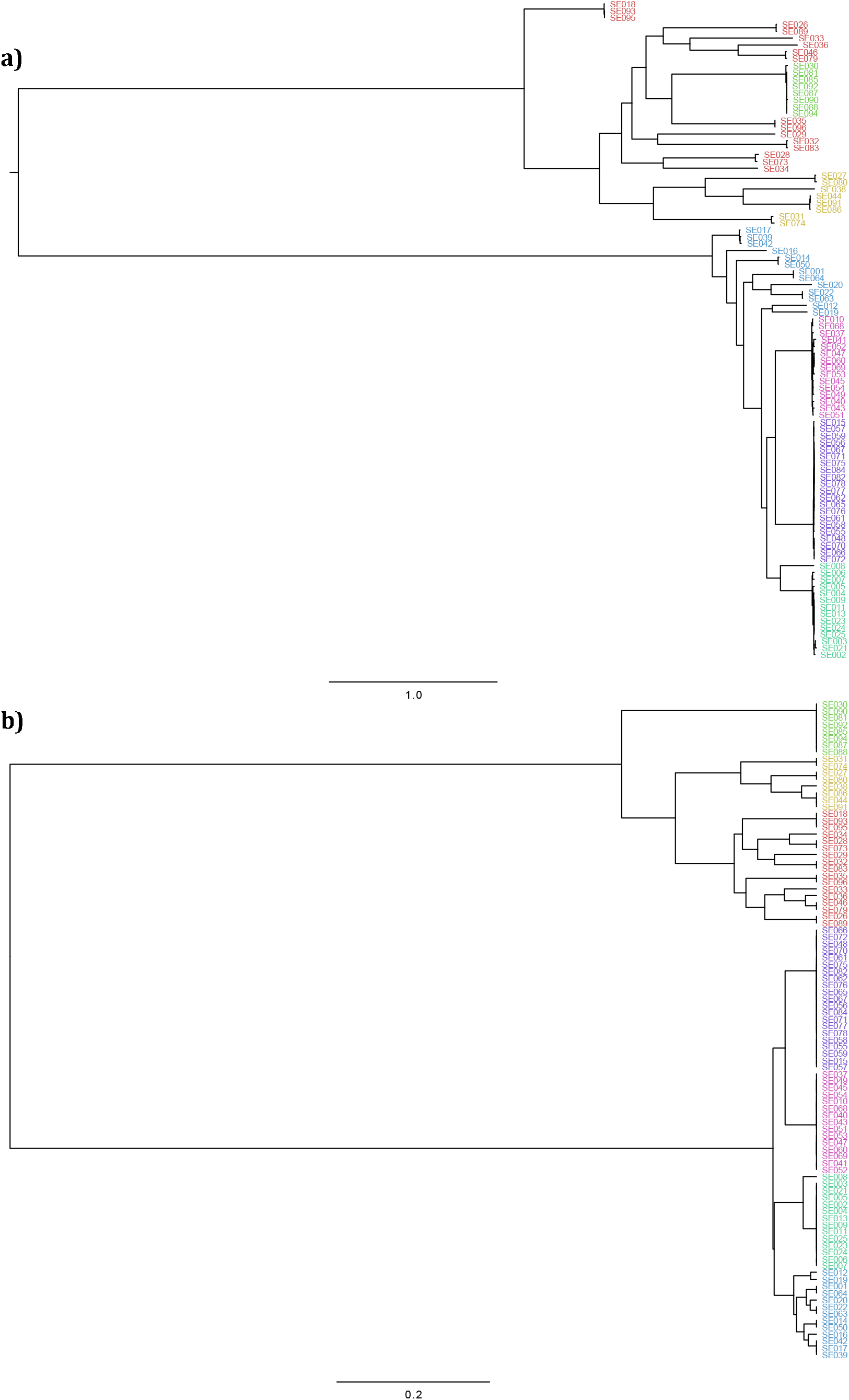
a) Tree used for Monte Carlo simulations 733 ions of 96 *S. pneumoniae* genomes. b) UPGMA tree from k-mer distance matrix produced from simulated reads. Colours are hireBAPS clusters.

**Supplementary figure 3:**
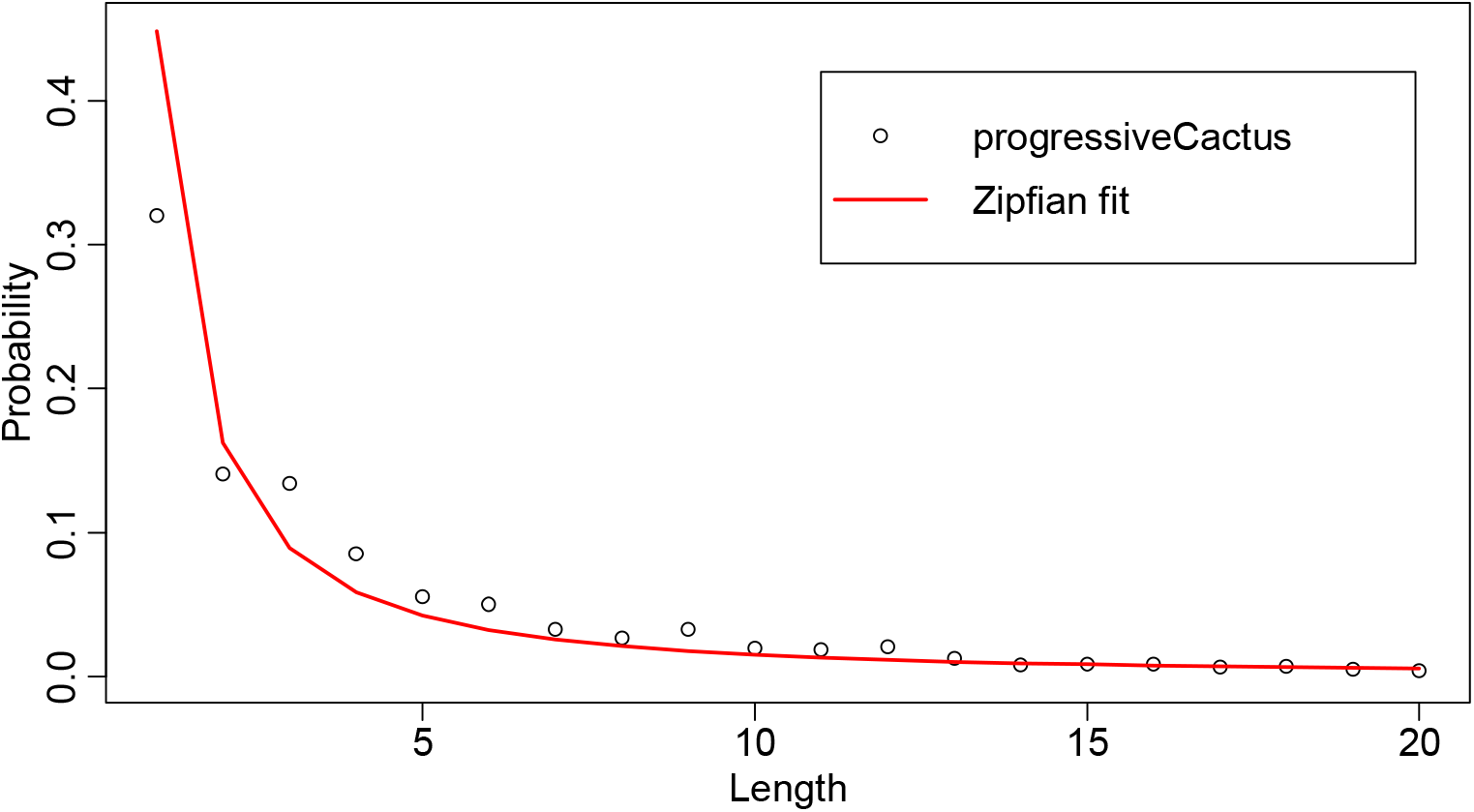
Estimated size distribution for INDELs, as estimated from a Progressive Cactus alignment of three members of the *Streptococcus* genus. A power law *p=L^k^* (Zipfian function; *p* is probability, *L* is INDEL length, *k* is a free parameter) is fit to the data, the parameter *k* is used in the simulations.

**Supplementary figure 4:**
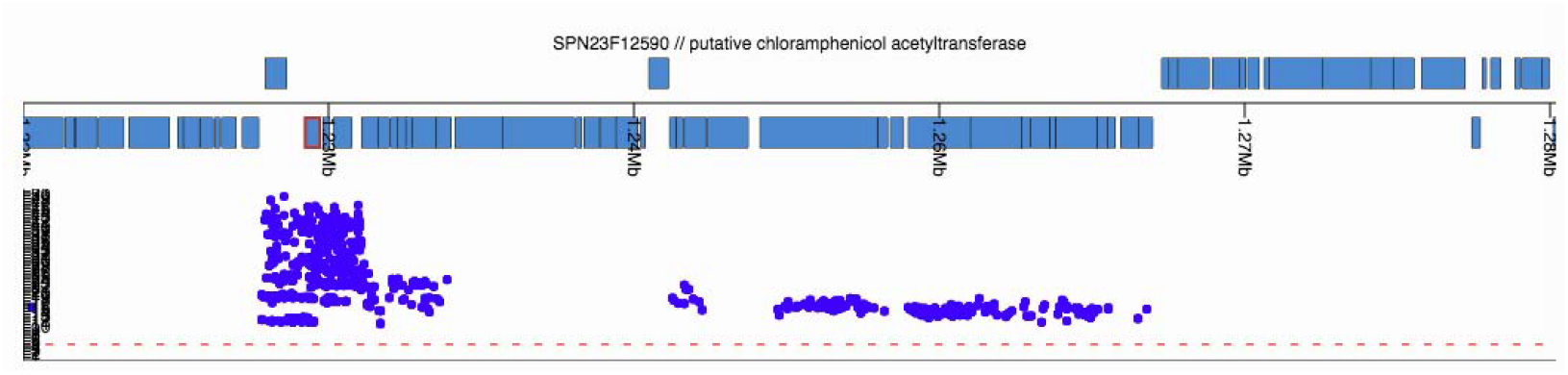
JScandy view of ATCC 700669 reference genome (blue blocks at top genes on forward and reverse strands) and Manhattan plot of start positions of the 1 508 of 1 526 k-mers significantly associated with chloramphenicol resistance which map to the integrative conjugative element (ICE) Tn*5253*. The hits are all in within the ICE, and the most significant hits cluster around the *cat* gene (which is outlined in red).

**Supplementary figure 5:**
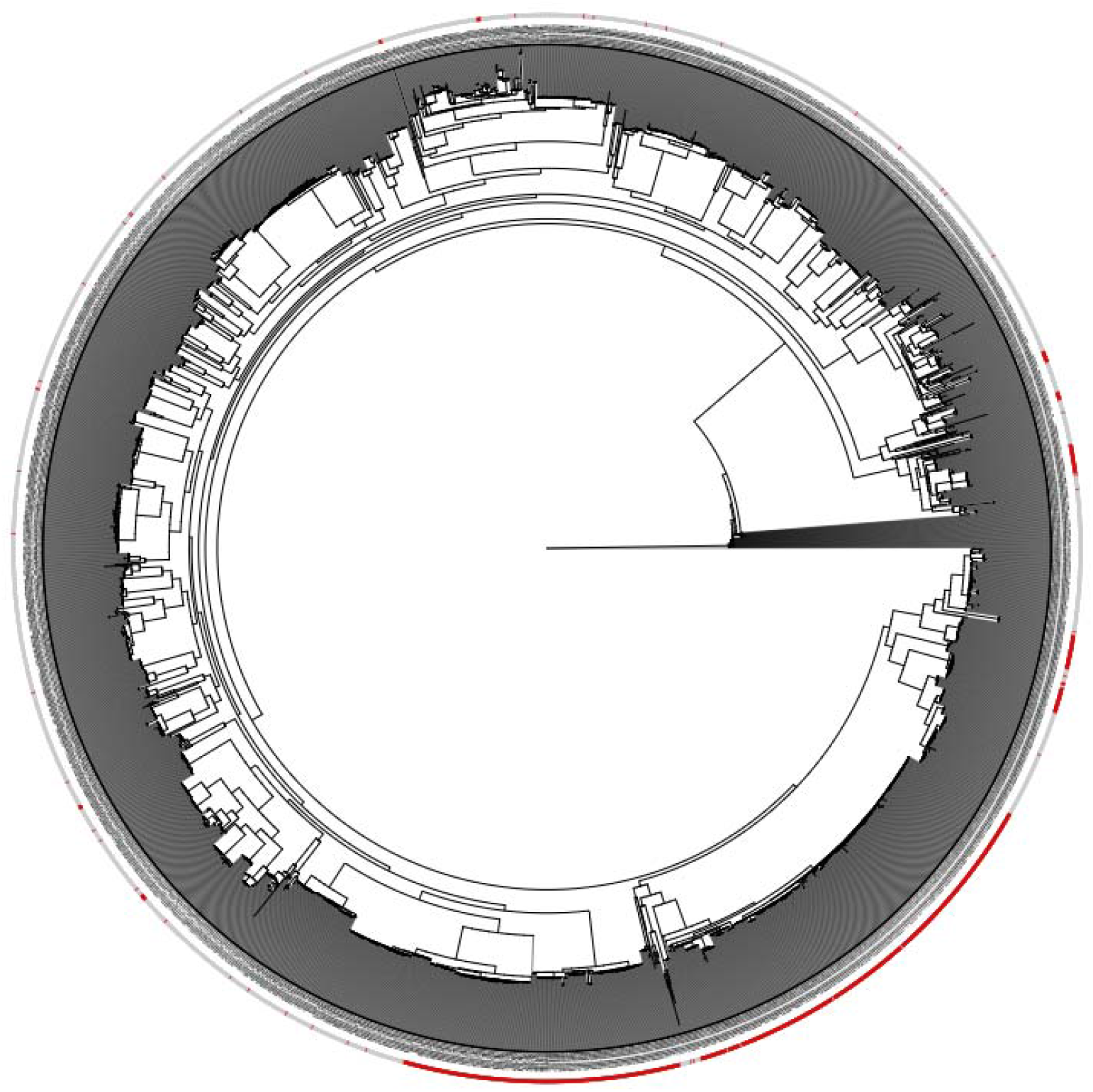
Neighbour joining tree from Chewapreecha *et. al*. study of 3069 Thai carriage *S. pneumoniae* samples, from a SNP alignment produced by mapping to the ATCC 700669 reference strain. Outer ring: red if resistant to Erythromycin, grey if sensitive.

**Supplementary figure 6:**
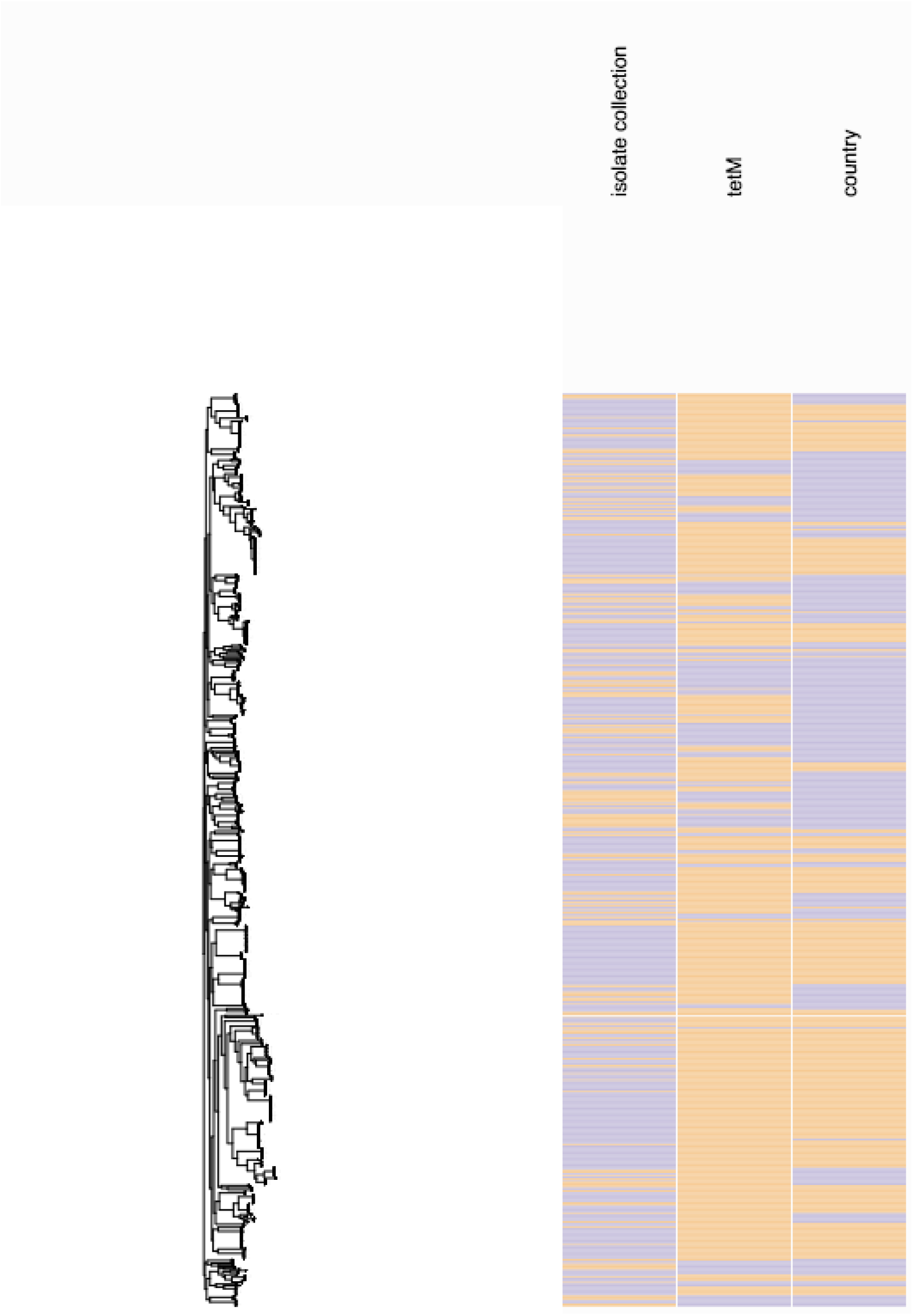
JSCandy view of S. pyogenes metadata on the right, showing whether isolates are invasive/non-invasive (orange/purple), presence of *tetM* (orange – absent, purple – present) and country of isolation (orange – Fiji, purple – Kilifi). Tree from a core genome alignment of all isolates is drawn on the left, with tips aligned to the metadata.

**Supplementary figure 7:**
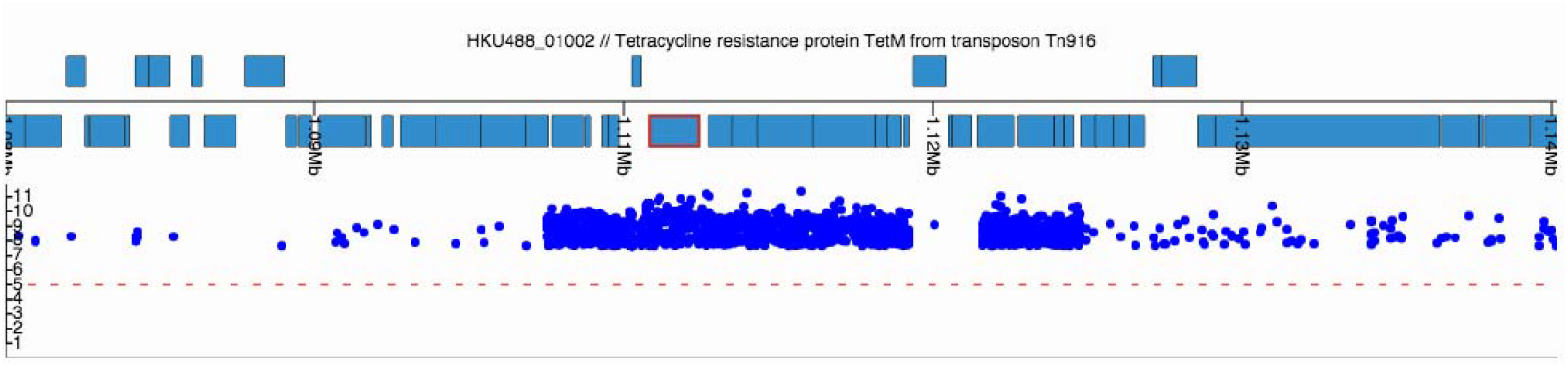
JScandy view of *S. pyogenes* HKU488 reference genome (blue blocks at top genes on forward and reverse strands, *tetM* highlighted in red) and Manhattan plot of start positions of k-mers significantly associated with invasiveness when not adjusted for country of origin.

**Supplementary figure 8:**
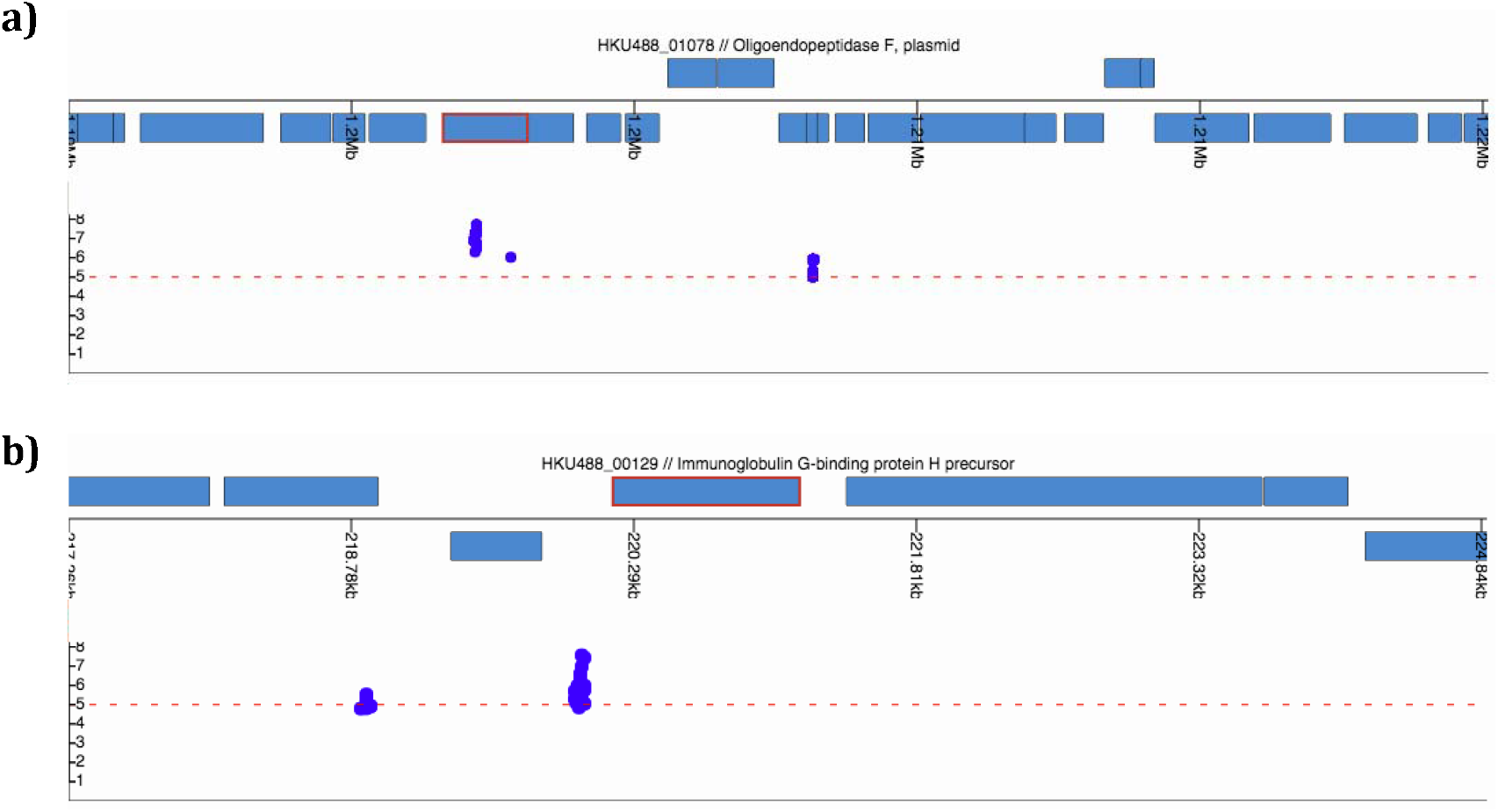
As supplementary figure 7, except with the Manhattan plot showing p-values when adjusted for country of isolation. a) *pepF*; b) IgG binding protein H precursor.

**Supplementary figure 9:**
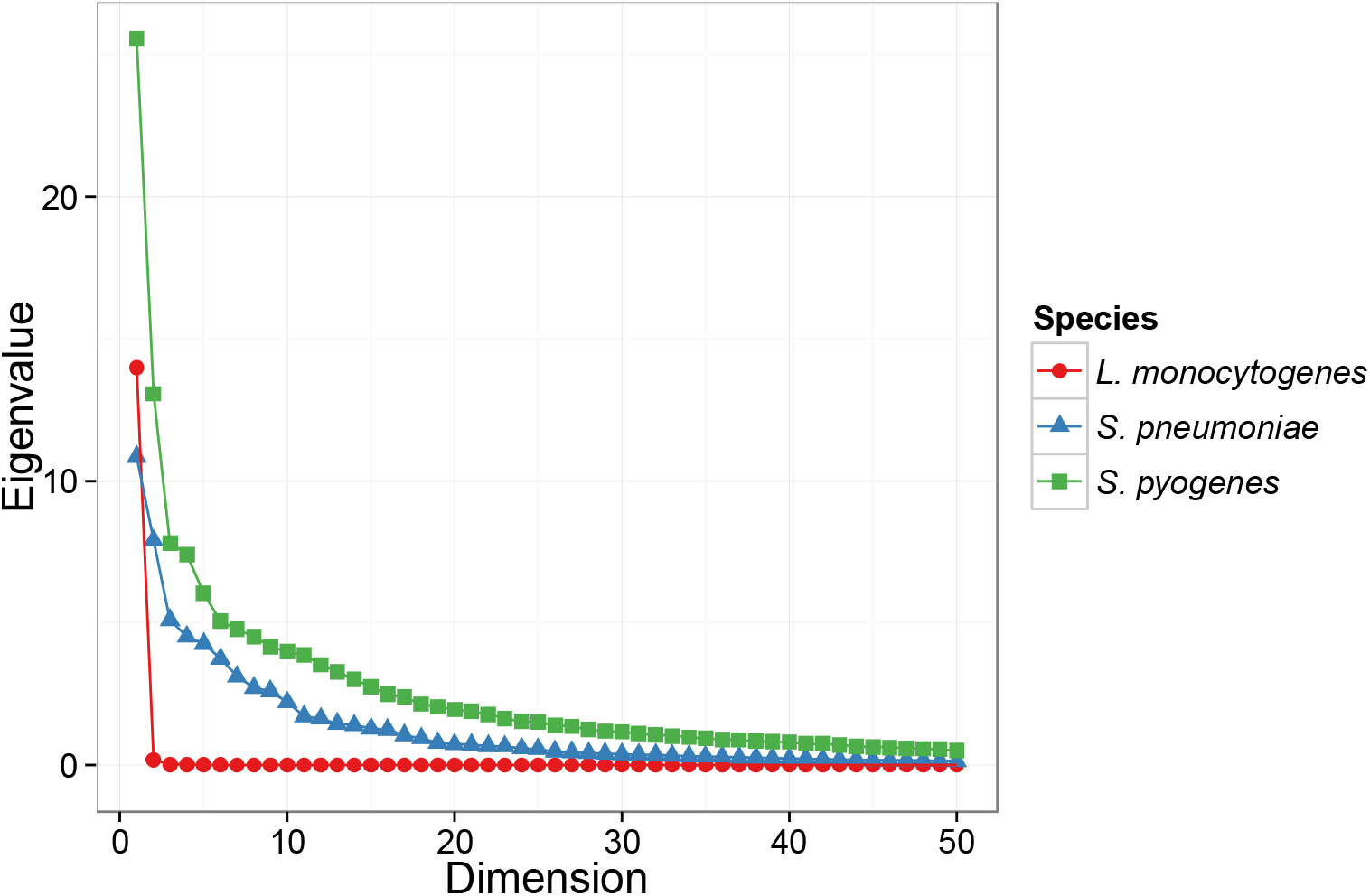
Scree plot for the first fifty dimensions of the 96 *Listeria monocytogenes* isolates (Supplementary figure 2) in red, 3 069 *Streptococcus pneumoniae* isolates (Supplementary figure 5) in blue, and 675 *Streptococcus pyogenes* isolates (Supplementary figures 6 and 7) in green.

